# Social stress causes gut dysbiosis in male, female, and aggressor mice

**DOI:** 10.1101/2025.06.26.661806

**Authors:** Isabel Garcia, Fatmanur Kilic, Chris-Ann Bryan, Jose Castro-Vildosola, Sai Anusha Jonnalagadda, Akhila Kasturi, Jaqueline Tilly, Jordyn Smith, Sofia Valentin, Sergio Moncayo, Tamara Hala, Eric Klein, Brian F. Corbet

## Abstract

**BACKGROUND:** Psychological stress causes gut dysbiosis, which is associated with adverse effects on physical and mental health in humans and mice. Understanding the specific taxa of gut bacteria changed by stress, and whether stress differentially alters their relative abundance in males and females, has important implications for stress-related disorders.

**METHODS:** We modeled individual differences in resilience or susceptibility using the chronic social defeats stress (CSDS) paradigm. Here, C57BL/6 mice are exposed to a novel retired breeder CD-1 aggressor for 10 minutes per day for 10 days. In this paradigm, resilient and susceptible subpopulations can be identified using the social interaction paradigm following CSDS. Fecal samples were collected following Day 1 and Day 10 of CSDS. 16S ribosomal RNA sequencing was used to identify the relative abundance of 200 bacteria species. We analyzed group differences in phyla, genera, and species in CD-1 aggressors along with resilient, susceptible, and non-stressed control male and female C57/BL/6 intruders.

**RESULTS:** Stress reduced microbiome diversity and caused gut dysbiosis in all groups, including aggressors. CSDS altered the relative abundance of every phylum detected. We report genera whose relative abundance was changed by CSDS or sex. Increases in the relative abundance of an uncultured *Ruminococcus* species on day 1 predicted aspects of social behavior, with a stronger correlation in stressed females compared to males.

**CONCLUSIONS:** CSDS causes gut dysbiosis in male and female mice, with generally similar effects in mice that behave differently following stress.

**ONE SENTENCE SUMMARY:** We report the effects of chronic social defeat stress on gut dysbiosis in aggressors, non-defeated controls, and male and female mice that are either resilient or susceptible to the adverse effects of stress.

**IN THIS ISSUE SUMMARY:** We characterize changes to the gut microbiome of male and female mice that are either resilient or susceptible to the adverse effects of social stress. Stress shifted the relative abundance of broad and specific taxa, which sometimes differed between males and females. We found that the abundance of an uncultured *Ruminococcus* species on day 1 predicted social behavior following stress. This effect was primarily observed in females.

## INTRODUCTION

Psychological stress is associated with disturbance of the gut microbiome, a complex and dynamic community of microorganisms. Gut dysbiosis, which refers to disturbances in the relative abundance of taxa in the gut microbiome, has been reported in Major Depressive Disorder (MDD)^1^, generalized anxiety disorder^2^, and posttraumatic stress disorder (PTSD)^3,4^. Inflammatory bowel diseases characterized by gut dysbiosis, like Chron’s disease, share comorbidity with symptoms of psychological stress^4–8^. Symptoms of depression or anxiety are associated with more severe symptoms of inflammatory bowel disease in the future and inflammatory bowel disease diagnosis is associated with the development of depression or anxiety symptoms^5^. Therefore, there is a reciprocal relationship between inflammatory diseases of the gut with stress-related psychiatric disorders.

The Chronic Social Defeat Stress (CSDS) paradigm is a useful model for understanding the relationships among stress, gut dysbiosis, and stress-related behavior^9^. Following 10 days of social defeat stress from a CD-1 aggressor, social interaction testing reveals distinct resilient and susceptible subpopulations of C57BL/6 mice. Susceptible mice display less time in a social interaction zone with a target mouse present compared to absent. They also display anxiety- and anhedonia-like behavior^10^ along with changes in brain circuits^11,12^, glucocorticoid levels^13^, and brain inflammation^14^. Resilient mice, which spend more time in a social interaction zone with a target mouse present, behave similar to non-stressed, co-housed controls.

Rodent behavior is influenced by the microbiome as fecal transplants from humans with MDD^15^ or rats susceptible to the adverse effects of stress^16^ cause certain maladaptive behavioral changes in non-stressed rodents. The gut microbiome has been investigated in resilient and susceptible male mice following CSDS^17–19^, although gut dysbiosis following CSDS in females has not been confirmed as far as we are aware. Sex differences in stress-induced gut dysbiosis might underlie the observation that certain inflammatory bowel diseases are generally more prevalent and severe in women compared to men in Western countries^20–24^. Inflammatory bowel diseases contribute to symptoms of depression and anxiety^5^, so sex differences in gut dysbiosis might also contribute to the higher prevalence of stress-related psychiatric disorders in women compared to men^25^. Additionally, to the best of our knowledge, the effects of CSDS on gut dysbiosis in aggressor mice is not known. Characterizing gut dysbiosis in aggressors, in addition to controls, is important for determining whether taxa altered in stressed C57BL/6 mice are altered because of stress or transfer from the CD-1 aggressor.

Here, we use 16S ribosomal ribonucleic acid (rRNA) to quantify the relative abundance of 200 bacteria species in fecal samples. We investigated the effects of Day (Day 1 and Day 10), Sex (male and female), and Stress group (aggressors, non-defeated, resilient, and susceptible). We found that CD-1 aggressors had increased alpha diversity compared to stressed groups. Beta diversity, which measures an overall difference in bacteria species, was altered by CSDS, sex, and stress groups. The abundance of every detectable phylum was altered by CSDS. *Firmicutes*, *Proteobacteria, Verrucomicrobiota*, *Cyanobacteria*, and *Desulfobacteria* were also different in C57BL/6 males compared to females. We also report each genus changed by CSDS or sex within each subgroup. We did not detect differences in genera between resilient and susceptible mice, but we found that Day 1 abundance of uncultured species from the *Oscillobacter*, *Ruminococcus*, and *Negativibacillus* genera predict aspects of social behavior following CSDS. Together, our findings characterize changes in the abundance of gut microbiome taxa following CSDS, how this differs between males and females, and which species best predict behavior following CSDS.

## METHODS AND MATERIALS

### Mice

10 to 14-week-old male and female C57BL/6 mice were used. Littermates were matched between non-defeated and stressed groups and cohoused until the first day of CSDS. For aggressors, retired breeder male CD-1 mice were purchased from Charles Rivers Laboratories (Wilmington, MA) and singly housed for two weeks prior to the beginning of CSDS. Mice were kept on a 12-12 light-dark cycle (7:00 – 19:00). All Experiments were performed in compliance with all relevant ethical regulations for animal testing and research. Experiment protocols followed the NIH Guide for the Care and Use of Laboratory Animals and were approved by the Rutgers University Institutional Animal Care and Use Committee.

### Chronic Social Defeat Stress (CSDS) and fecal sample collection

Mice were subjected to CSDS for 10 consecutive days as in^9,13,26^. For CSDS, C57BL/6 mice (intruders) are placed in the home cage of a CD-1 aggressor. These home cages are clear, polysulfone hamster cages that are 26.7 cm (w) × 48.3 cm (d) × 15.2 cm (h) (Allentown, cat. no. PC10196HT). Mice physically interact for 10 minutes. For females, 10 µL of urine from a CD-1 aggressor was applied to the tail immediately prior to daily physical interaction as in^26^. Intruders and aggressors are cohoused, but separated using a custom, clear polysulfone partition with 54 perforated holes (0.3 cm in diameter, Allentown), allowing aggressor CD-1 mice and intruder C57BL/6 to see, smell, and hear each other but not physically interact. Every 24 hours, intruders are placed in the cage of a novel aggressor. Non-defeated controls are cohoused in the same type of partitioned cage, but never physically interact. This is repeated for 10 days in total. All bedding and feces were removed from the cage immediately following CSDS, just before mice were partitioned on Day 1 and Day 10. Fecal samples from were collected within 60 minutes following partitioning on Day 1 and Day 10 and frozen at -80C until metagenome sequencing.

### Social interaction paradigm

Social interaction was performed 24 hours following the tenth day of CSDS. Mice were placed in a 30 cm x 38 cm social interaction chamber with 9 cm x 9 cm corner zones. The social interaction zone is a 25.5 cm x 10 cm rectangle with an arc that reaches out to an additional 3.75 cm at its peak. Target mice are placed in an upside down, metal pencil holder with perforated sides that are 7.5 cm in diameter. Mice exposed to CSDS were placed in the social interaction box for 150 seconds without the target mouse, removed for 30 seconds while a CD-1 target mouse is placed in its enclosure, and returned to the social interaction box for an additional 150 seconds. All behavior in the social interaction chamber without and with the target mouse is video recorded. Time in either corner and time in the social interaction zone were scored by two observers blind to the experimental groups. 90% scoring precision was confirmed, and means were calculated. Social interaction was conducted in red light. Social interaction ratios were calculated as time in the social interaction zone with the target mouse present/absent. K-means clustering analysis was used to identify the social interaction ratio value that best separates resilient compared to susceptible groups and found to be 1.255.

### 16S rRNA sequencing

Fecal samples were shipped on dry ice to SeqCenter (Pittsburgh, PA) for sample preparation and metagenome sequencing. DNA was extracted using the ZymBIOMICS DNA miniprep kit (Zymo Research, cat. no. D4304). DNA libraries were prepared used a *Quick*-16s Plus Next Generation Sequencing Library Prep kit (Zymo Research, cat. no. D6421) to selectively amplify the hypervariable V3/V4 region of the gene coding 16S rRNA. V3/V4 amplicons were sequenced at 100k read depth using an Illumina platform. Sequences were denoised using the Qiime2 dada2 plugin. Sequence reads were aligned to bacteria species genomes using the Silva-138 database. The number of sequence reads that aligned to each bacteria species was used to determine their relative abundance.

### Statistical Analyses

RStudio was used for all analyses. Data were subjected to the Shapiro-Wilks test for normal distributions and Bartlett’s test for heterogeneity. Only social interaction ratio and the relative abundance of *Bacteroidetes* passed both assumptions required for General Linear Models and were analyzed by two- and three-way analysis of variance (ANOVA), respectively. For non-parametric testing, Aligned Rank Transform (ART) ANOVA was used to detect main effects (Day, Sex, and Stress group) and their interactions. For post hoc differences that included aggressors, estimated marginal means with Bejamin-Hochbrerg p-values correction was used. Differences between groups different by one factor (e.g. resilient males on Day 1 compared to resilient females on Day 1) are reported. To identify differences in the abundance of each genus between subgroups, α was adjusted to 0.01. The Shannon-Weiner Index and Bray-Curtis tests were used to assess alpha and beta diversity, respectively. To identify correlations between the relative abundance of species and behavior, Spearman’s Rank Correlation was used and α was adjusted to 0.001. Fisher’s r-to-z transformation was used to identify differences in correlation strength in males compared to females. All code and resulting statistics are available at https://github.com/bcorbett24/CSDS-microbiome-250611/blob/main/RStudio%20code.

## RESULTS

### Social behavior in male and female mice following CSDS

Social interaction (SI) testing was performed 24 hours following a tenth consecutive day of CSDS (Figure 1A). A significant stress group effect was observed for SI ratios. This was primarily driven by males as all male stress groups behaved differently from one another. SI ratios were higher in resilient males compared to non-defeated (ND) and susceptible males and lower in susceptible males compared to ND males. A trend for social interaction ratios being higher in resilient females compared to susceptible females was observed (Figure1B). Females spent more time in the social interaction zone than males (Supplemental Figure 1). Across all groups, females spent less time in the corner compared to males. The presence of a target mouse generally reduced time in spent in the corner across all groups. Non-defeated and resilient males displayed reduced time in the corner with a target mouse present compared to absent. A trend for reduced time in the corner with the target mouse present compared to absent was observed for resilient females (Figure 1C). These findings characterize sex differences in the behavioral phenotypes defining resilient and susceptible subpopulations.

**Figure 1.**
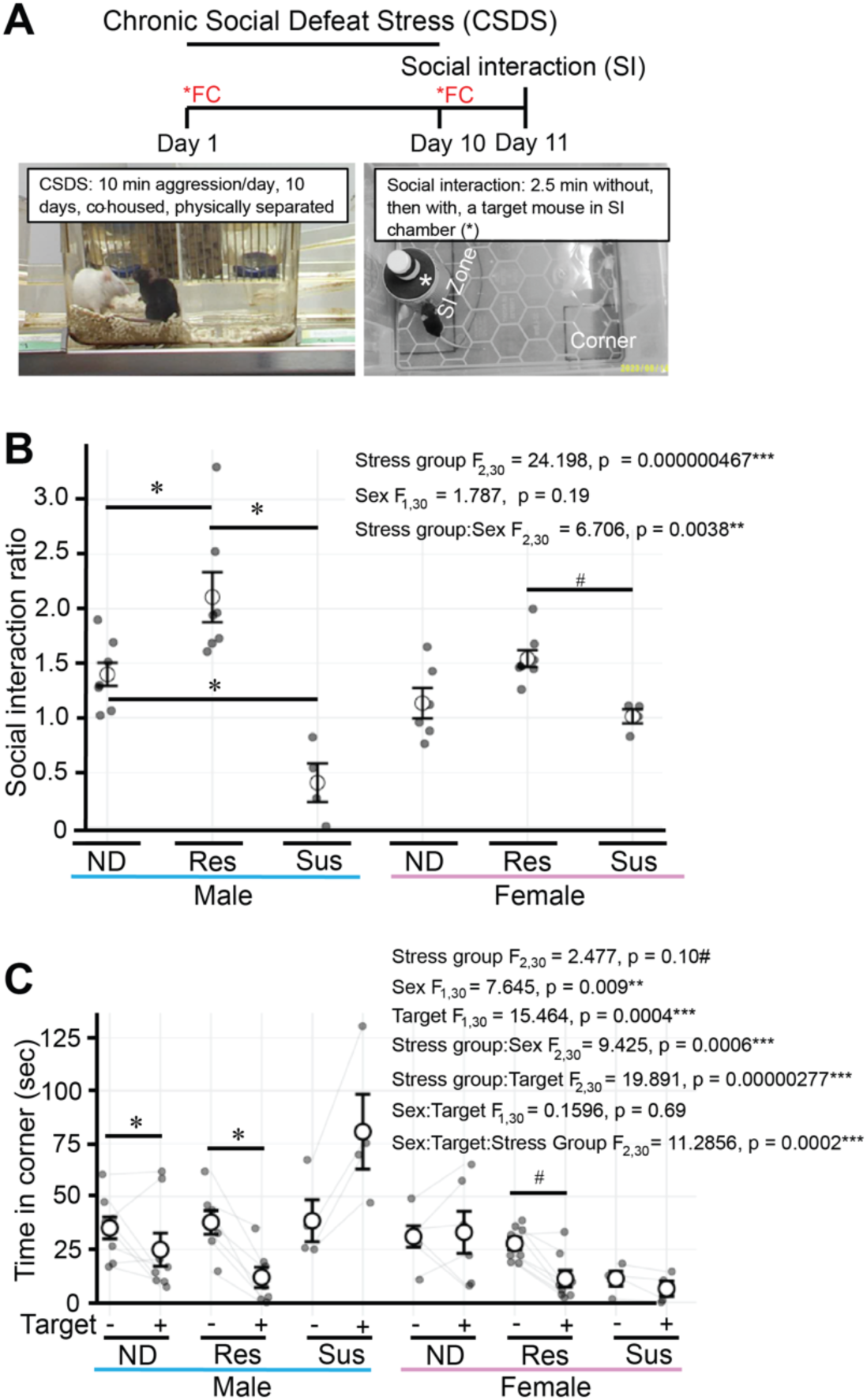
Effects of Chronic Social Defeat Stress in males and females**. A)** Schematic of experimental timeline demonstrating fecal sample collection (FC) immediately following Days 1 and 10 of CSDS. The social interaction (SI) paradigm was performed on Day 11. **B)** Compared to non-defeated (ND) controls, SI ratio was increased in resilient (RES) males and decreased in susceptible (SUS) males. A trend for reduced SI ratio in resilient compared to susceptible females was observed. Two-way ANOVA with estimated marginal means post-hoc. **C)** ND and Res males spent less time in the corner with a target mouse present. A trend for reduced time in the corner with a target mouse present was observed for resilient females. Paired Aligned Rank Transform (ART) ANOVA with estimated marginal means post-hoc. Ends of horizontal lines represent post-hoc comparisons. Bars represent mean ± SEM. * adjusted p < 0.05, # adjusted p < 0.10. Males: ND n = 8, Res n = 7, Sus n = 4; Females: ND n = 6, Res n = 7, Sus n = 4.

### Alpha diversity is higher in CD-1 aggressors compared to C57BL/6 intruders

16S ribosomal RNA sequencing was used to quantify the relative abundance of gut bacteria species (relative abundance of each species for each mouse in all groups is provided in Supplemental Table 1). The Shannon Index, which accounts for species abundance and the evenness of their distribution^16,27,28^, was used to compare alpha diversity of the gut microbiome within each group. An interaction between Day and Stress group was observed. Post hoc testing revealed that alpha diversity was higher in aggressors compared to ND males, but not resilient or susceptible males, on Day 1. Shannon Index scores were reduced on Day 10 compared to Day 1 in all male and female groups exposed CSDS except susceptible males, which trended toward a decrease. On Day 10, Shannon Index scores were higher in aggressors compared to all other male groups. Alpha diversity was not changed in non-defeated males or females. Sex differences in Shannon Index were not observed within any stress group on any day. Shannon Index scores were higher in resilient and susceptible females compared to non-stressed control females on Day 1, but not Day 10 (Figure 2). Together, these findings indicate that CSDS reduces alpha diversity is higher in CD-1 aggressors compared to C57BL/6 males and reduced by CSDS.

**Figure 2.**
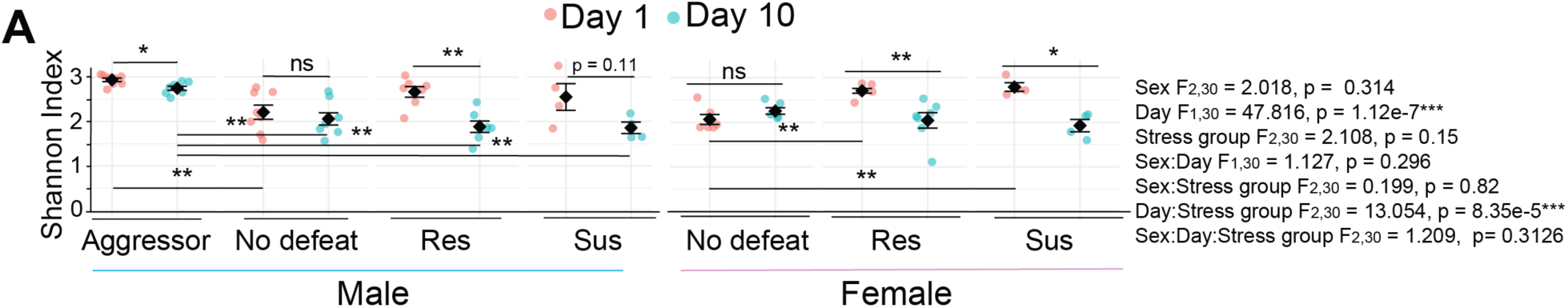
Effects of CSDS on alpha diversity. **A)** Shannon index values are higher in aggressors compared to intruders and higher in Res and Sus males and females on Day 1 compared to Day 10. Effects and their interactions assessed using Paired Aligned Rank Transform (ART) ANOVA with estimated marginal means post-hoc with Benjamini-Hochberg used to correct for p-values. Bars represent mean ± SEM. Ends of horizontal lines represent post-hoc comparisons. * adjusted p < 0.05, ** adjusted p < 0.01. Males: Aggressors n = 9, No defeat n = 8, Resilient (Res) n = 7, Susceptible (Sus) n = 4; Females: No defeat n = 6, Res n = 7, Sus n = 4.

### Bray-Curtis distance is increased by stress and sex

Bray-Curtis distance, which represents beta diversity, measures shifts in the relative abundance of species among groups. Day, Sex, and Stress group effects were observed along with all interactions. Male subgroups were statistically different from their female counterpart with the only exception being susceptible mice on Day 1. Most stress groups were different from one another within each sex on either day. The only exceptions were that resilient males were not different from non-defeated or susceptible males on Day 1 or susceptible males on Day 10, and resilient females were not different from susceptible females on Day 1 or non-defeated and susceptible females on Day 10. Bray-Curtis distance was different between non-defeated females and susceptible, but not resilient, females on Day 10. However, Bray-Curtis distance is not different between resilient and susceptible females on Day 10 (Figure 3A, Table 1). To better visualize beta diversity induced by CSDS, we graphed aggressor, non-defeated, resilient, and susceptible mice separately. Regarding Day effects, Bray-Curtis distance was different on Day 1 and Day 10 in all male and female groups except non-defeated males and females (Figure 3B). Together, these findings indicate that sex, CSDS, and mouse breed contribute to beta diversity.

**Figure 3.**
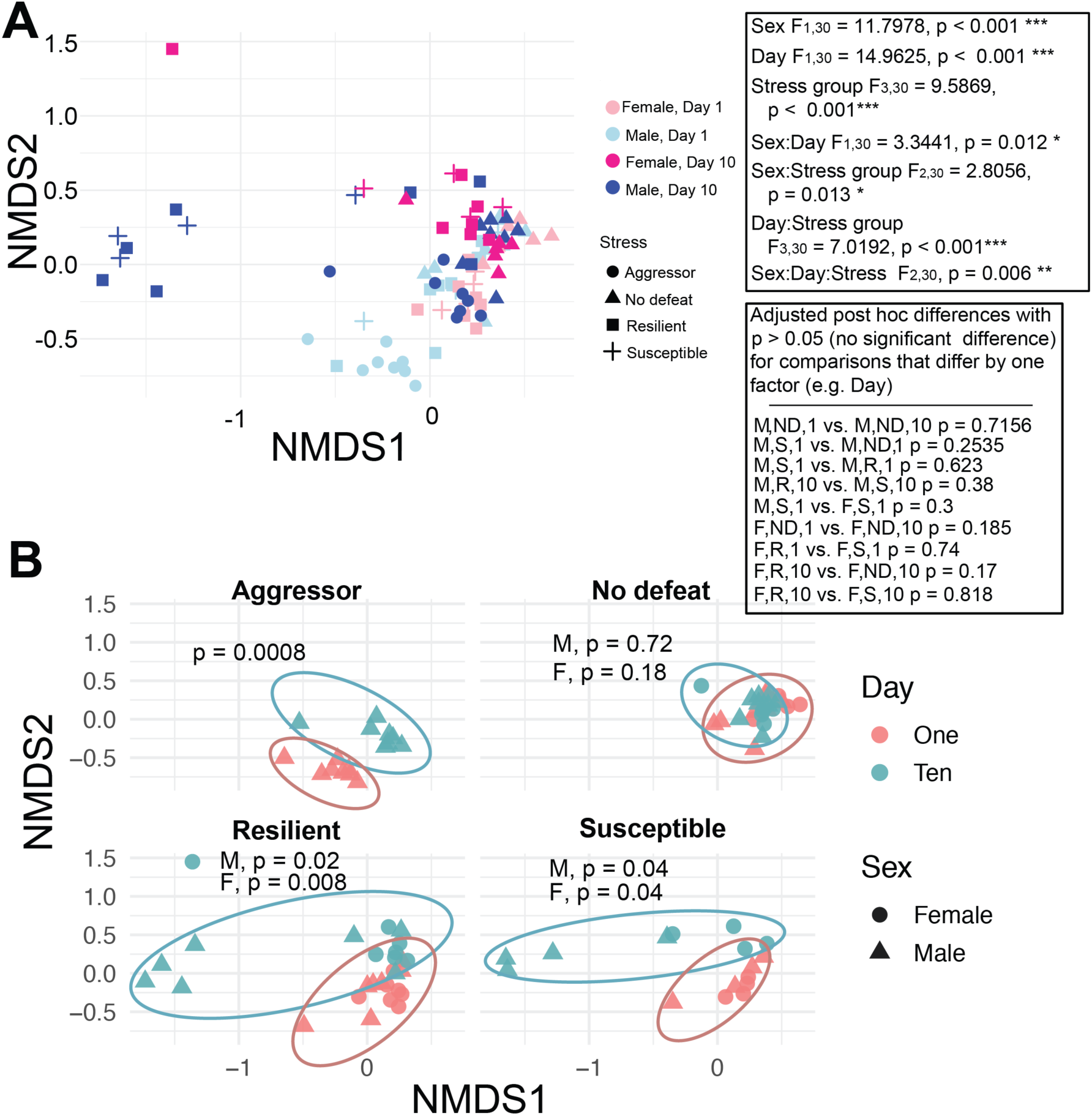
Effects of CSDS on beta diversity. **A)** Plots non-parametric multidimensional scaling (NMDS) coordinates 1 and 2 for each mouse following Bray-Curtis analysis of species data. Effects and their interactions assessed using PERMANOVA. Pairwise PERMANOVA using Bonferroni false discovery rate for p-adjustment was used to identify differences between subgroups. Most subgroups were significantly different from one another, only pairwise subgroup differences with p < 0.05 are shown. All pairwise comparisons are available in Table 1. **B)** Data are shown on separate graphs for each Stress group to highlight Bray-Curtis distance between Days 1 and 10 was similar for non-defeated males and females, but it was different for all groups exposed to CSDS. Eclipses fitted to capture 90% of data within each group. * p < 0.05, ** p < 0.01, *** p < 0.0001. Males: Aggressors n = 9, No defeat n = 8, Resilient (Res) n = 7, Susceptible (Sus) n = 4; Females: No defeat n = 6, Res n = 7, Sus n = 4.

**Table 1:**
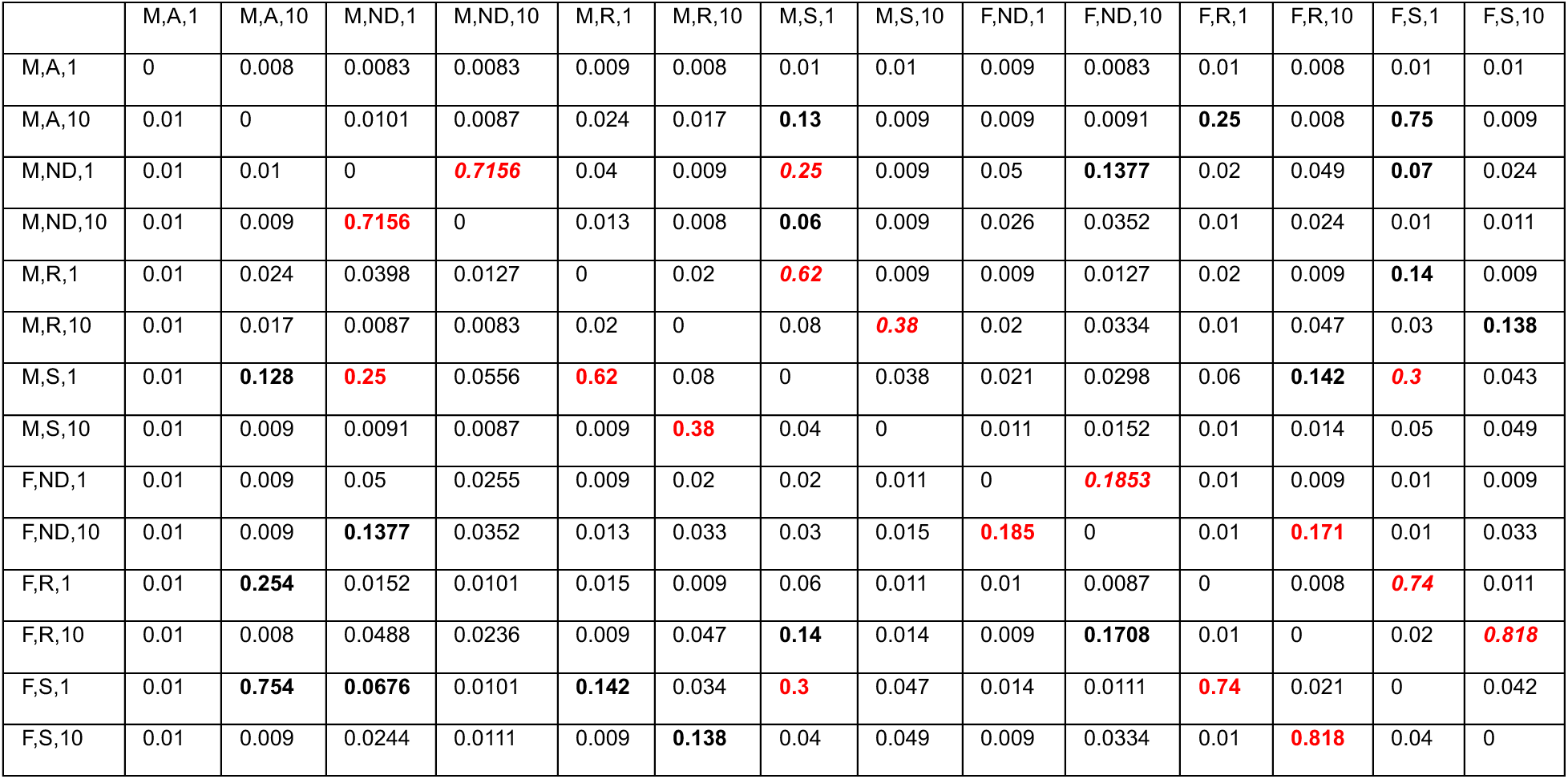
Bray-Curtis analysis post hoc differences. The adjusted p-values derived from pairwise PERMANOVA comparisons between the groups represented in the first row and first column. All p-values > 0.05 are bolded. Red p-values represent differences between two groups that differ by only one factor. Cells with a repeated value, with the identical comparison shown to the left, are Italicized. M, male; F, female; A, aggressor; ND, non-defeated control; R, resilient; S, susceptible; 1, Day one; 10, Day ten.

### Effects of sex and stress on *Firmicutes* and *Bacteroidetes*

Bray-Curtis distances indicate overall shifts in the relative abundance of gut microbiome taxa, but identification of differences in specific taxa requires direct investigation. *Firmicutes* and *Bacteroidetes* are the most abundant bacteria phyla in the gut of mice^27^ and humans^28^. Effects of Day and Stress group were observed. A Day x Stress group interaction and post-hoc testing found reduced *Firmicutes* abundance was attributed to aggressors, resilient, and susceptible mice but not non-defeated controls. An overall Sex effect was observed for higher *Firmicutes* in females compared to males. On Day 10, resilient and susceptible males and females displayed lower *Firmicutes* abundance compared to aggressors and their same-sex non-defeated control (Figure 4A). A significant overall Day effect indicated that 10 days of CSDS increases the abundance of *Bacteroidetes*. A Day x Stress group interaction and post-hoc testing revealed this Day effect was driven by stress as significant increases were observed in aggressors along with resilient or susceptible, but not non-defeated control, males and females (Figure 4B). A reduced ratio in the abundance of *Firmicutes* to *Bacteroidetes* (FB ratio) is a biomarker for Chron’s disease and ulcerative colitis^29^. A significant overall Day effect indicated that 10 days of CSDS reduced the FB ratio. A Day x Stress group interaction and post-hoc testing revealed this was driven by CSDS as significant reductions were observed in aggressors along with resilient or susceptible, but not non-defeated control, males and females. Together, these findings indicate that CSDS alters the abundance of the two most abundant bacteria phyla in the gut in males and females.

**Figure 4.**
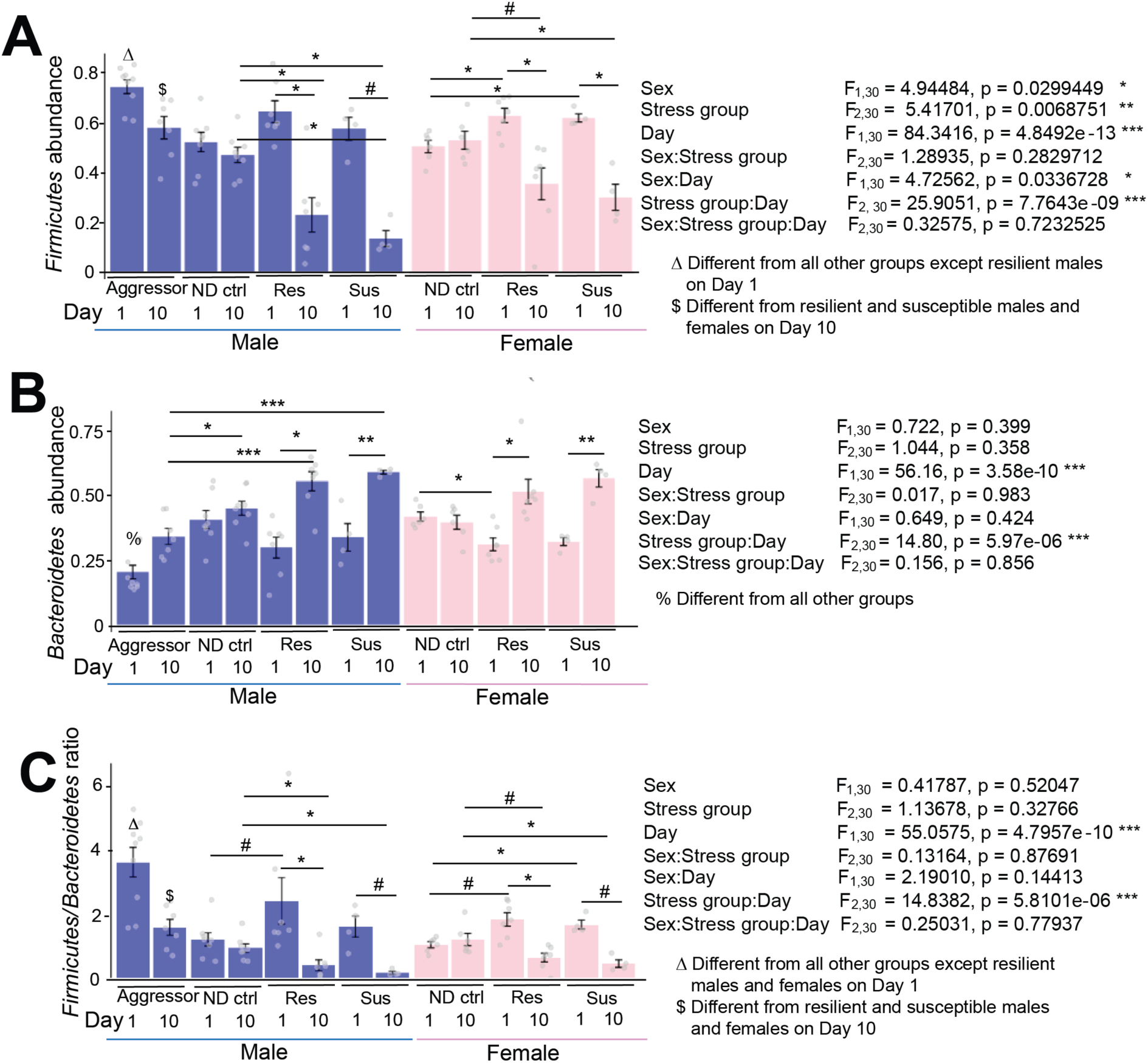
CSDS reduces *Firmicutes* and increases *Bacteroidetes* in males and females. **A)** The relative abundance of *Firmicutes* is higher in aggressors compared to all groups other than resilient males on Day 1, higher in aggressors compared to resilient and susceptible males on Day 10, generally higher in females compared to males across all groups (Sex effect), and reduced in males and females exposed to CSDS, but not non-defeated controls, on Day 10 compared to Day 1. **B)** The relative abundance of *Firmicutes* is lower in aggressors compared to all other groups on Day 1, lower in aggressors compared to other male groups on Day 10, and increased in males and females exposed to CSDS, but not changed in non-defeated controls, on Day 10 compared to Day 1. **C)** The ratio of relative abundance of *Firmicutes/Bacteroidetes* is higher in aggressors compared to all groups other than resilient males on Day 1, higher in aggressors compared to resilient and susceptible males and females on Day 10, and reduced in males and females exposed to CSDS, but not non-defeated controls, on Day 10 compared to Day 1. For A and C, main effects and their interactions were assessed using Aligned Rank Transform (ART) ANOVA with estimated marginal means post-hoc. For B, main effects and their interactions were identified using three-way ANOVA since assumptions of the general linear model were met. Estimated marginal means post hoc testing with Benjamini-Hochberg correction of p-values was used to identify differences between subgroups. Bars represent mean ± SEM. Ends of horizontal lines represent post-hoc comparisons. * adjusted p < 0.05, ** adjusted p < 0.01, *** adjusted p < 0.0001, # p < 0.10. Males: Aggressors n = 9, No defeat (ND) n = 8, Resilient (Res) n = 7, Susceptible (Sus) n = 4; Females: No defeat n = 6, Res n = 7, Sus n = 4.

### Effects of sex and stress on less abundant gut bacteria phyla

Day, Sex, and Stress group effects, along with their interactions, were observed for *Proteobacteria* and *Verrucomicrobiota*. These effects were primarily driven by resilient and susceptible males as *Proteobacteria* abundance was greater and *Verrucomicrobiota* trended towards an increase in resilient and susceptible males, but not females, on Day 10 compared to Day 1. No Day effects in *Proteobacteria* or *Verrucomicrobiota* were observed in aggressors or non-defeated controls (Figure 5A,B). *Actinobacteria* abundance was reduced by CSDS as indicated by a Day x Stress group interaction. Post hoc testing revealed a significant reduction in resilient females on Day 10 compared to Day 1. Compared to resilient females, susceptible female displayed a greater abundance of *Actinobacteria* on Day 1. Susceptible males displayed lower *Actinobacteria* compared to non-defeated males on Day 10. Aggressors displayed greater abundance of *Actinobacteria* compared to resilient and susceptible males and females on Day 10 (Figure 5C). *Cyanobacteria* and *Deferribacterota* abundance were reduced by CSDS and higher in males compared to females as indicated by overall Day, Sex, and Stress group effects along with interactions of these factors. Compared to non-defeated males and females, generally higher abundance of these phyla in was observed in aggressors on Days 1 and 10 and resilient and susceptible mice on Day 1, but not Day1 10, (Supplemental Figure 2 A,B).

**Figure 5.**
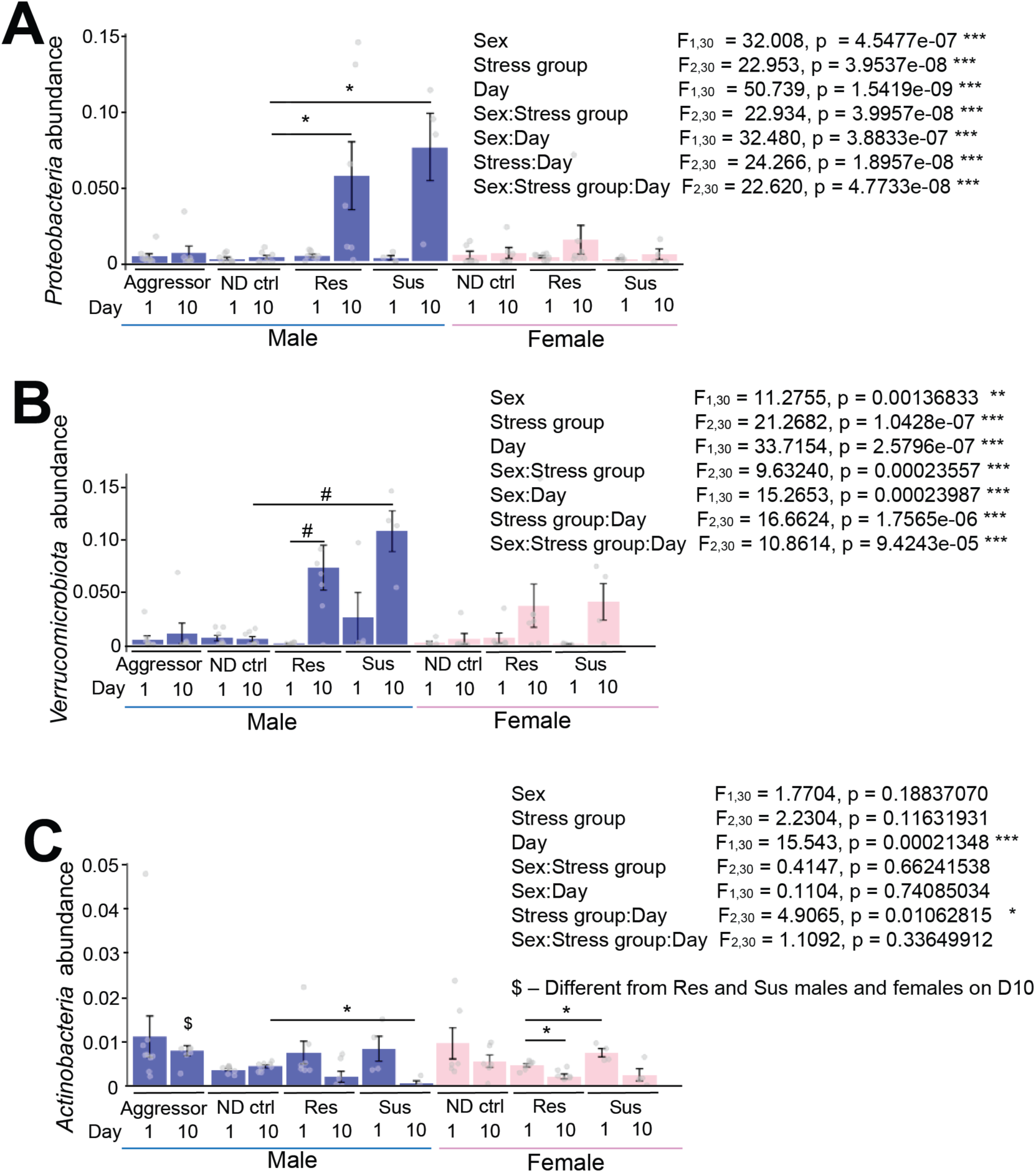
Effects of CSDS and sex on less abundant gut bacteria phyla. **A)** *Proteobacteria* abundance is increased in resilient and susceptible males, but no other groups, on Day 10 compared to Day 1. **B)** A trend for increased *Verrucomicrobiota* abundance is observed in resilient males on Day 10 compared to Day 1 and in susceptible males compared to non-defeated males on Day 10. **C)** *Actinobacteria* abundance is higher in aggressors compared to resilient and susceptible males and females on Day 10, lower in susceptible males compared to non-defeated males on Day 10, and reduced on Day 10 compared to Day 1 in resilient females. Main effects and their interactions were assessed using Aligned Rank Transform (ART) ANOVA with estimated marginal means post hoc testing using Benjamini-Hochberg correction of p-values to identify differences between subgroups. Bars represent mean ± SEM. Ends of horizontal lines represent post-hoc comparisons. * adjusted p < 0.05, # p < 0.10. Males: Aggressors n = 9, No defeat (ND) n = 8, Resilient (Res) n = 7, Susceptible (Sus) n = 4; Females: No defeat n = 6, Res n = 7, Sus n = 4.

*Desulfobacteria* abundance was minimal or completely absent although resilient females had higher abundance on Day 1 compared to all other groups on either day (Supplemental Figure 2C). Together, these results indicate that the relative abundance of *Proteobacteria* and *Verrucomicrobiota* are increased by CSDS, especially in males. The abundance of *Actinobacteria* and *Deferribacterota* in resilient and susceptible males on Day 1, but not Day 10, was similar to the abundance of these phyla from aggressors on Day 1. This might be attributed to rapid transfer of these less abundant phyla from aggressors to intruders.

### Effects of CSDS on the relative abundance of gut bacteria genera

The relative abundance of 23 genera, 20 of which being *Firmicutes*, were different on Day 10 compared to Day 1 in at least one male or female stress group. Except *Allistepes*, which was increased in aggressors, all other differences in the relative abundance of genera were observed in resilient or susceptible males or females. Other than *Paludicoloa*, which was increased in resilient males, all differentially abundant genera on Day 10 compared to Day 1 in males or females were reduced. *Lachnospiraceae*, *Butyricicoccus*, *Ruminococcus*, and an uncultured genus from Family XIII were reduced in both males and females. *Ruminococcus* was reduced in susceptible, but not resilient, females. Every other genus reduced by CSDS in the *Firmicutes* phylum was sex-specific, although trends for reduced abundance of that genus were sometimes observed for the other sex (Table 2, means and SEM provided in Supplemental Table 2). Together, these findings demonstrate that CSDS reduces genera in the *Firmicutes* phylum in males and females, whereas the relative abundance of genera in CD-1 aggressors and non-defeated controls were stable. No differences in the relative abundance of any genus were detected when comparing resilient compared to susceptible males and females.

**Table 2.**
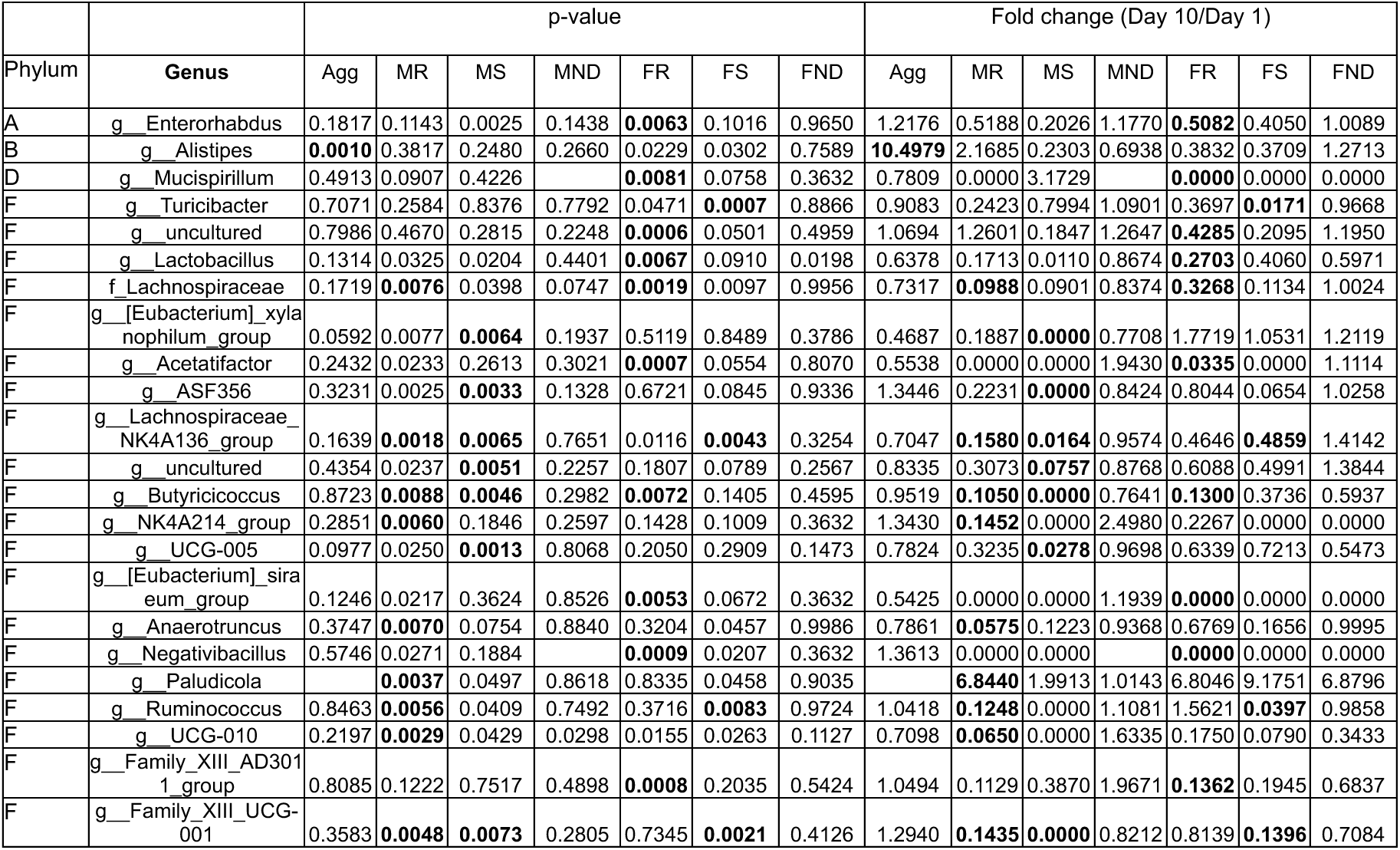
CSDS reduces *Firmicutes* genera in males and females. Differentially abundant genera on Day 1 compared to Day 10 in CD-1 aggressors (Agg), and C57BL/6 male (M) and female (F) resilient (R), susceptible (S), and non-defeated control (ND) mice. The left side of the table displays p-values following a paired t-test to compare Day 1 and Day 10. Comparisons in which p < 0.01 are bolded. The right side of the table displays fold change of Day 10 abundance divided by Day 1 abundance. A, *Actinobacteria*; B, *Bacteroidetes*; D, *Deferribacterota*; F, *Firmicutes*.

### Differences in the relative abundance of gut bacteria genera in males and females

On Day 1, *Lactobacillus* was higher in non-defeated females compared to non-defeated males. Trends for higher *Lactobacillus* were observed in females compared to males in all other Stress groups on Day 10. On Day 10, non-defeated females displayed reduced abundance of two uncultured genera, including the uncultured genus from Family XIII that was also reduced by CSDS, compared to non-defeated males. On Day 10, susceptible females displayed greater abundance of *Adlercreutzia* and lower abundance of *Bacteroides* compared to susceptible males. Seven of the 14 genera differentially abundant between males and females were observed in resilient mice on Day 1, but not Day 10. (Table 3, means and SEM provided in Supplemental Table 3). Together, these results indicate that *Lactobacillus* is higher in the gut of females compared to males, but most other genera are similarly abundant.

**Table 3.** Sex differences in the abundance of gut microbiome genera. Differentially abundant genera in females (F) compared to males (M) on Day 1 and Day 10 in resilient (R), susceptible (S), and non-defeated control (ND) mice. The left side of the table displays p-values comparing males and females. Comparisons in which p < 0.01 (unpaired t-test) are bolded. The right side of the table displays fold change in the relative abundance of females divided by males in that group on that day. Empty cells indicate that abundance of that genus was below the threshold for detection in males and female within that stress group. A, *Actinobacteria*; B, *Bacteroidetes*; F, *Firmicutes*.

| Phylum | Genus | p-value |  |  |  |  |  | Fold change (F/M) |  |  |  |  |  |
| --- | --- | --- | --- | --- | --- | --- | --- | --- | --- | --- | --- | --- | --- |
|  |  | D1ND | D10ND | D1R | D10R | D1S | D10S | D1ND | D10ND | D1R | D10R | D1S | D10S |
| A | **g_Adlercreutzia** | 0.1808 | 0.0853 | 0.2983 | 0.6328 | 0.0620 | **0.0064** | 0.3006 | 0.2844 | 1.5111 | 1.4509 | 4.5623 | **M=0** |
| B | **g_Bacteroides** | 0.4087 | 0.1339 | 0.4310 | 0.0135 | 0.7491 | **0.0049** | 0.4528 | 0.1710 | 0.7067 | 0.0194 | 0.7603 | **0.1155** |
| B | **g_Alistipes** | 0.3262 | 0.7080 | **0.0001** | 0.8012 | 0.0038 | 0.1301 | 0.3944 | 0.7227 | **6.6444** | 1.1741 | 6.4746 | 10.426 |
| F | **g_Holdemania** | 0.2643 |  |  |  | **0.0038** |  |  |  |  |  | **M=0** |  |
| F | **g_Turicibacter** | 0.0330 | 0.0281 | 0.1173 | 0.2375 | **0.0027** | 0.2777 | 2.0267 | 1.7974 | 2.2258 | 3.3958 | **3.2753** | 0.0700 |
| F | **g_Lactobacillus** | **0.0018** | 0.0244 | 0.0355 | 0.0370 | 0.4632 | 0.0245 | **2.4081** | 1.6577 | 2.2348 | 3.5265 | 1.3675 | 50.556 |
| F | **g_Clostridia_UCG-014** | 0.1715 | 0.1672 | **0.0026** | 0.0749 | 0.3415 | 0.6047 | 0.4518 | 0.7184 | **0.4918** | 0.2775 | 0.7471 | 0.5926 |
| F | **g_Lachnospiraceae_FCS020_group** | 0.0108 | 0.6428 | **0.0018** | 0.5464 | 0.1750 | 0.1490 | 0.4300 | 0.8559 | **0.3434** | 1.5388 | 0.6021 | 4.3737 |
| F | **g_Lachnospiraceae_NK4A136_group** | 0.0314 | 0.1004 | **0.0068** | 0.2505 | 0.6597 | 0.0242 | 0.4028 | 0.5950 | **0.7059** | 2.0761 | 0.9035 | 26.721 |
| F | **g_Monoglobus** | 0.9549 | 0.5947 | **0.0060** | 0.0520 | 0.0327 |  | 0.9670 | 0.7257 | **2.3983** |  | 2.5346 |  |
| F | **g_Oscillibacter** | 0.1267 | 0.4390 | **0.0005** | 0.1100 | 0.0289 | 0.7452 | 0.6461 | 1.1692 | **0.5123** | 2.2792 | 0.5917 | 1.3383 |
| F | **g_UCG-010** | 0.6245 | **0.0049** | 0.8732 | 0.4358 | 0.6625 | 0.3559 | 1.2087 | **0.2540** | 1.0386 | 2.7964 | 1.1728 |  |
| F | **g_Anaerovorax** |  | 0.2199 | **0.0059** | 0.7752 |  | 0.5653 |  |  | **0.0000** | 1.5183 |  | 1.8744 |
| F | **g_Family_XIII_UCG-001** | 0.5768 | **0.0083** | 0.3308 | 0.0359 | 0.2342 | 0.3559 | 0.7900 | **0.6815** | 0.8227 | 4.6644 | 1.4354 |  |

### Correlations between the abundance of gut bacteria species with behavior

On Day 1 of CSDS, the abundance of three gut bacteria species in the *Oscillospirales* order of the *Clostridia* class correlated with subsequent social interaction behavior following CSDS. An uncultured species in the *Oscillobacter* genus negatively correlated with time in the social interaction zone with a target mouse present (Figure 6A). An uncultured species in the *Ruminococcus* genus negatively correlated with time in the social interaction zone (Figure 6B) and positively correlated with time in either corner when a target mouse was present, with a stronger correlation in females compared to males (Figure 6C). An uncultured species in the *Negativibacillus* genus, also in the *Ruminococcaceae* family, negatively correlated with time in either corner when a target mouse was present (Figure 6D). This genus was not detectable on Day 10. No correlations between social behavior and the ratio of species abundance on Day 10 divided by abundance on Day 1 were observed. No correlations with p < 0.0001 were observed when comparing behavior and the relative abundance of each species on Day 10. Together, these results demonstrate that the abundance of bacteria species in the *Clostridia* class on Day 1 correlate with subsequent social behavior. *Ruminococcus* abundance on Day 1 correlates particularly well with social anxiety-like behavior in females. A full list of all species correlating with social interaction ratio, time in the social interaction zone, or time in the corner with or without a target mouse present are presented in Supplemental Table 4.

**Figure 6.**
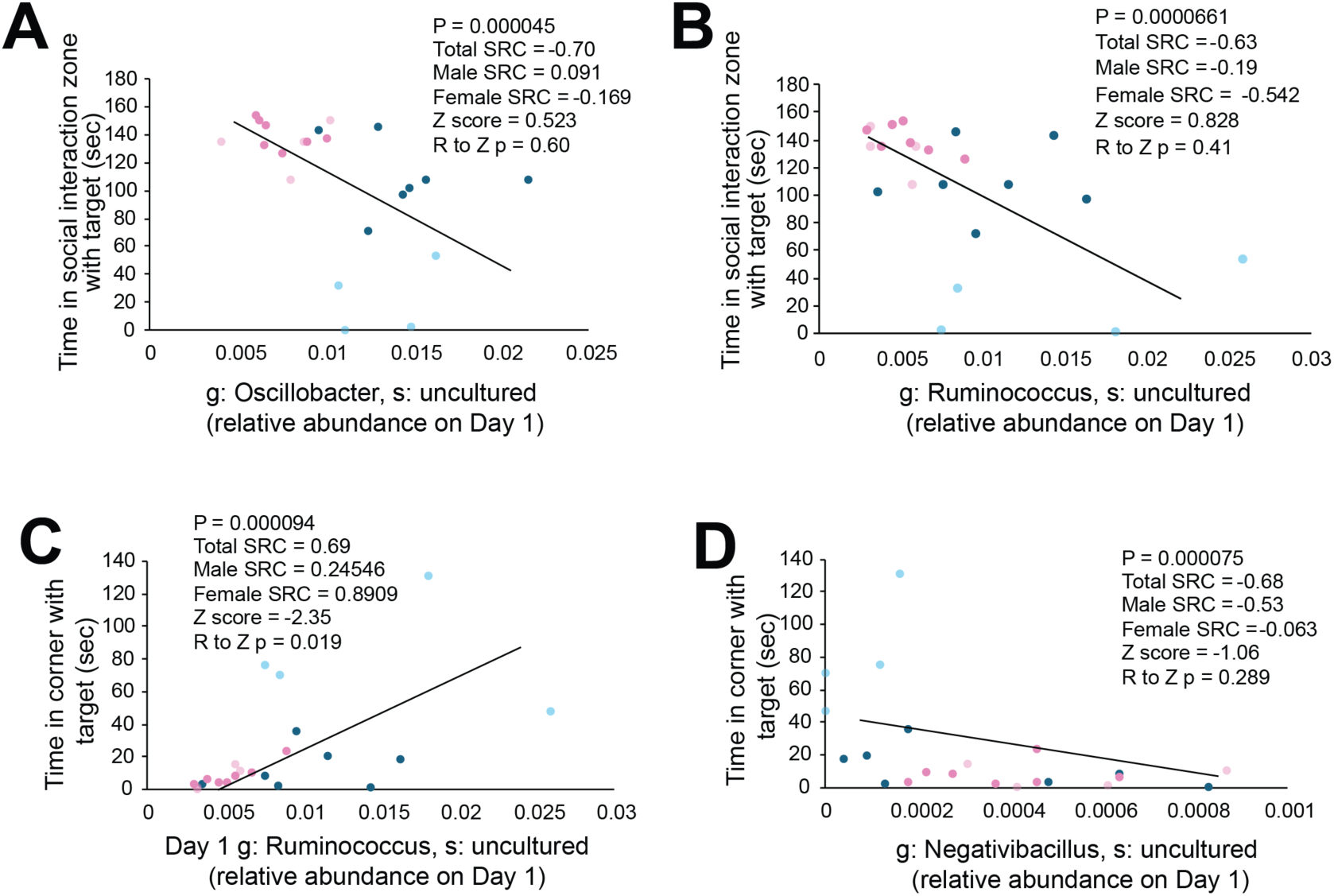
The abundance of uncultured species in the *Clostridia* class predict social behavior following stress. Combining males and females, the abundance of uncultured species (s) in the **A)** *Oscillobacter* genus (g) and **B)** *Ruminococcus* genus on Day 1 negatively correlate with time in the social interaction zone when the target mouse is present. **C)** Abundance of the same uncultured *Ruminococcus* species correlates with time in the corner when the target mouse is present, and this correlation is stronger in females compared to males. **D)** Combining males and females, the relative abundance of an uncultured species in the *Negativibacillus* genus negatively correlates with time in the corner when the target mouse in present. P-values are derived from Spearman rank correlation (SRC) using males and females combined. R-to-Z p-values are provided to identify differences in the correlation strength in males compared to females.

## Discussion

Here, we characterize gut dysbiosis caused by CSDS and how this differs among resilient and susceptible males, females, and aggressors. Our finding that stress reduces alpha diversity in males and females is consistent with studies in humans^30^ and male mice^31^, although effects in females were not known. On Day 1, susceptible females displayed higher alpha diversity compared to non-defeated females. This was likely due to transfer from aggressors, which displayed higher alpha diversity compared to intruders. Despite an initial increase in alpha diversity of Day 1, alpha diversity was reduced following 10 days of CSDS in all stressed groups but was not lower than alpha diversity of non-defeated controls on Day 10. We collected samples following Day 1 of CSDS to account for transferrable species from aggressors, which were presumably also present on Day 10. Although transfer from aggressors was a likely contributor to increased alpha diversity on Day 1, a single social defeat causes gut dysbiosis compared to baseline microbiome composition^32^. Microbiota composition changes throughout the gastrointestinal tract^33^, so acute effects caused by stress might rapidly mobilize species of gut bacteria not normally detectable in feces. Beta diversity was altered by Day, Sex, and Stress effects along with their interactions. Post hoc testing of Bray-Curtis distances revealed differences in most subgroups. For both males and females, beta diversity was similar in resilient and susceptible mice, so CSDS causes similar effects on beta diversity regardless of behavior following CSDS.

We found that CSDS alters the relative abundance of every phylum as assessed by Stress group:Day interactions. Post hoc testing revealed changes in the relative abundance of phyla in resilient and susceptible, but not non-defeated, mice. The effects of sex were more subtle, although there was an overall Sex effect or Sex:Stress group interaction for every phylum except *Bacteroidetes* and *Actinobacteria*, indicating sex influenced overall levels of most phyla and/or influenced how their relative abundance was changed by CSDS. We found that CSDS reduced *Firmicutes* and increased *Bacteroidetes,* thereby reducing the *Firmicutes/Bacteroidetes* ratio, which is a biomarker for Chron’s disease and ulcerative colitis^29^, two inflammatory bowel diseases that share comorbidity with symptoms of psychological stress^4–8^. *Proteobacteria* were increased by CSDS in males, but not females. A similar trend was observed for *Verrucomicrobiota*, but post hoc differences were not significant despite overall Day, Sex, and Stress group effects along with their interactions. *Actinobacteria* was reduced by CSDS, with significantly lower abundance in susceptible males compared to non-defeated males on Day 10. *Cyanobacteria* were generally elevated in males compared to females, although their abundance was still relatively low. Compared to non-defeated controls, *Deferribacterota* abundance was higher in aggressors, resilient, and susceptible mice on Day 1, suggesting transfer of this phylum from aggressor to intruder. *Desulfobacterota* was higher in resilient females on Day 1 compared to all other groups, but a high degree of variability was observed due to the relatively low abundance of this phylum. Gut dysbiosis of *Firmicutes*, *Bacteroidetes*, *Proteobacteria*, *Verrucomicrobiota*, and *Actinobacteria* are all observed in humans stress-related psychiatric disorders with the same directional changes^34–36^, indicating that stress-induced changes in the abundance of these phyla can be accurately modeled in mice using CSDS.

The majority of the genera altered by CSDS were members of the *Firmicutes* phylum and were reduced by CSDS. *Lachnospiraceae*, *Butyricicoccus*, *Ruminococcus* were reduced by CSDS in both males and females. *Butyricicoccus* and *Ruminococcus* are typically lower in humans with in stress-related psychiatric disorders^37,38^, whereas *Lachnospiraceae* abundance is less consistent^39^. There were no significant differences in the abundance of any genus between resilient and susceptible males or females. This might be at least partially because social interaction ratios, which dictate resilience and susceptibility, are less related to the abundance of genera than underlying aspects of behavior that influence social interaction ratio, like motivation, motility, or social anxiety-like behavior. The only significant correlations between species abundance and behavior correlated the abundance of species in the *Oscillospirales* order with time in the social interaction zone or time in the corner, but not social interaction ratio.

Abundance of *Oscillobacter*, *Negativibacillus,* and *Ruminococcus* on Day 1, but not Day 10 or the ratio of their abundance on Day 10 compared to Day 1, correlated with behavior. This might be because these species promote biological processes that increase susceptibility to the adverse effects of stress, like inflammation. Alternatively, abundance of these species could represent latent biomarkers indicative of vulnerability to the development of these processes occurring during CSDS or their abundance could be changed by stress-induced molecular or physiological mechanisms that also change behavior. *Oscillobacter* and *Ruminococcus* abundance negatively correlated with time in the social interaction zone with a target mouse present. *Ruminococcus* abundance also correlated with time in the corner with a target mouse present. *Oscillobacter* and *Ruminococcus* in the gut are decreased in MDD^40^, although increased *Oscillobacter* has also been reported^41^. *Ruminococcus* is also reduced in generalized anxiety disorder^42^. *Ruminococcus* might be a particularly good biomarker for stress-related disorders and/or a potential therapeutic target since its abundance is reduced by CSDS in males and females and humans with stress-related disorders. We also found its abundance is more strongly correlated with social avoidance in female mice compared to mice. This is importance since stress-related psychiatric disorders are generally twice as prevalent in women compared to men^25^.

Together, our findings demonstrate that CSDS causes gut dysbiosis in male and female mice. Whereas the relative abundance of every phylum was shifted by CSDS, only 23 of the 133 detectable bacteria genera were changed, indicating stress influences more general taxa than specific ones. Social exclusion^43^ and Generalized Anxiety Disorder^44^ are associated with reduced *Firmicutes* and increased *Bacteroidetes*, whereas the opposite is true for MDD^45–47^.

This might be because different physiological processes caused by stress, like activation of either the sympathetic nervous system or hypothalamic-pituitary-adrenal axis, induce gut dysbiosis. Future projects will investigate the physiological processes by which CSDS and other stress paradigms contribute to gut dysbiosis.

---

**Supplemental Figure 1.**
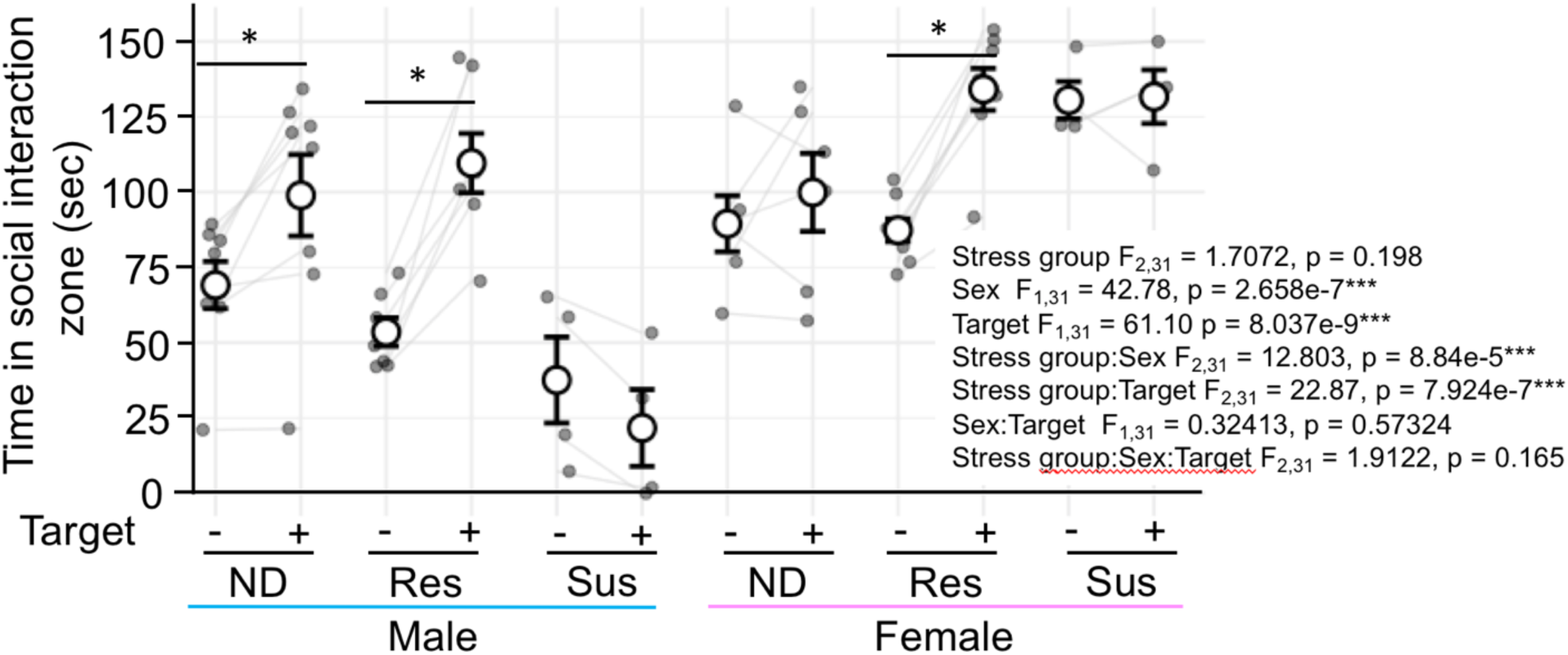
Time in social interaction zone in males and females following CSDS. Across all groups and target conditions, females spend more time in the social interaction zone. Non-defeated (ND) and resilient (Res), but not susceptible (Sus) males and females spend more time in the social interaction zone with the target mouse present. Repeated measures 3-way ANOVA with estimated marginal means post-hoc. Ends of horizontal lines represent post-hoc comparisons. Bars represent mean ± SEM. * adjusted p < 0.05. Males: ND n = 8, Res n = 7, Sus n = 4; Females: ND n = 6, Res n = 7, Sus n = 4.

**Supplemental Figure 2.**
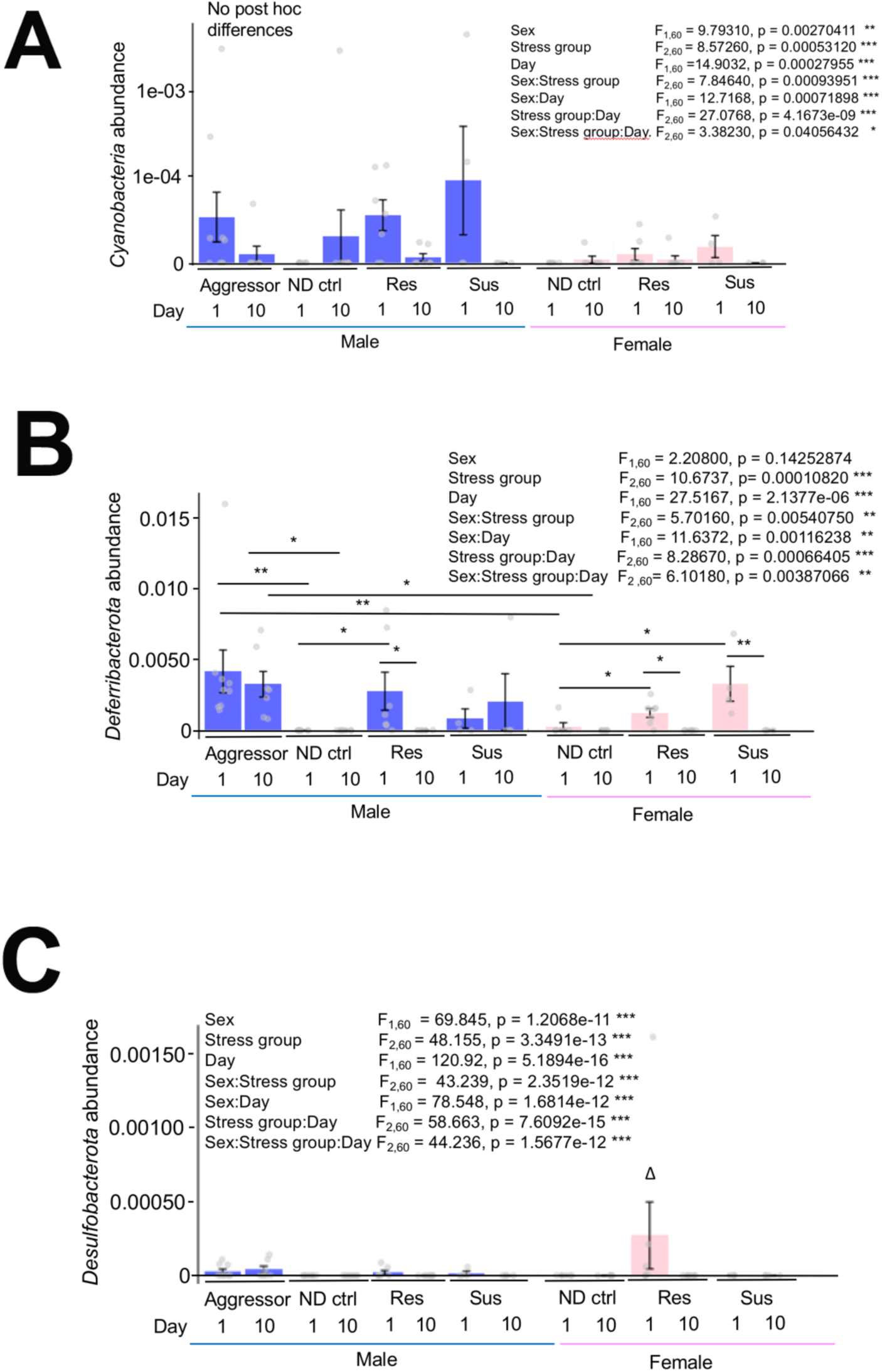
Effects of CSDS and sex on bacteria phyla with low abundance suggest transfer from aggressors . **A)** Across all groups, *Cyanobacteria* are higher in males compared to females. All main effects and their interactions are significant. **B)** All main effects and their interactions are significant for *Deferribacterota* other than sex. **C)** *Desulfobacterota* abundance is higher in resilient females on Day 1 compared to all other groups on Day 1 and Day 10, but this effect is primarily driven by one mouse. Main effects and their interactions were assessed using Aligned Rank Transform (ART) ANOVA with estimated marginal means post hoc testing using Benjamini-Hochberg correction of p-values to identify differences between subgroups. Bars represent mean ± SEM. Ends of horizontal lines represent post-hoc comparisons. * adjusted p < 0.05, # p < 0.10. Δ p < 0.05 for pairwise comparisons between every other group. Males: Aggressors n = 9, No defeat (ND) n = 8, Resilient (Res) n = 7, Susceptible (Sus) n = 4; Females: No defeat n = 6, Res n = 7, Sus n = 4.

